# Lifetime-based multiplexed detection of viral RNA using fluorogenic aptamers

**DOI:** 10.64898/2026.04.13.718069

**Authors:** Yuan-I Chen, Yu-An Kuo, Yujie He, Naseem Siraj, Emile J. Batchelder-Schwab, Yin-Jui Chang, Siem Yonas, Yuting Wu, Zhenglin Yang, Anh-Thu Nguyen, Sohyun Kim, Yi Lu, Chengde Mao, Pengyu Ren, Hsin-Chih Yeh

## Abstract

Fluorogenic aptamers (FAPs) are emerging molecular probes for viral RNA and DNA sensing. However, their use in multiplexed nucleic acid sensing has been hindered by cross-reactivity and overlapping emission spectra. Here we address these limitations by introducing a fluorescence-lifetime-based multiplexed detection strategy using variants of the DNA fluorogenic aptamer *Lettuce* that exhibits distinct fluorescence lifetimes when complexed with the fluorogen TO1-biotin. To effectively evolve *Lettuce* for diverse lifetimes, we developed a large-scale screening platform, termed FAP-FLIM-NGS (fluorogenic <u>ap</u>tamer-based fluorescence lifetime imaging <u>m</u>icroscopy on <u>n</u>ext-<u>g</u>eneration <u>s</u>equencing chips), which measures the fluorescence lifetimes of ∼10^4^ *Lettuce*/TO1-biotin complexes directly on an Illumina *MiSeq* flow cell. Using this approach, three variants with markedly different lifetimes were identified: a single mutant (smC14T, 6.0 ns) and two double mutants (dmA5T/C14T, 5.2 ns, and dmA5T/T22A, 4.4 ns). To demonstrate the utility of these *Lettuce* variants in multiplexed detection, a set of split *Lettuce* probes targeting viral RNA fragments derived from SARS-CoV-2, MERS-CoV, and influenza A were designed and tested. Phasor plot analysis confirmed that these probes can robustly distinguish individual targets as well as mixtures containing any two or all three targets purely based on distinct fluorescence lifetimes of probes, thereby overcoming the challenges of cross-reactivity and spectral overlap. Beyond this proof of concept, our findings establish a generalizable strategy for engineering FAPs with customized photophysical properties, opening new avenues for next-generation diagnostics and molecular sensing technologies.

## INTRODUCTION

Fluorescence lifetime measurement has added a new dimension to multiplexed fluorescence imaging and sensing^1-4^, which have traditionally relied on fluorescence color and emission intensity for target encoding or differentiation^5^. Although simultaneous measurement of fluorescence color and lifetime using a pulsed-interleaved excitation^6^ scheme can distinguish up to nine and six different tags/targets in fixed^1^ and live cells^4^, respectively, in a single scan, strategies to engineer tags for diverse lifetimes remain limited and inefficient. Current approaches often rely on structurally informed rational design^7, 8^, computational modeling^9^, or small-scale selection^10^ to create tags of different lifetimes. A notable example is the engineering of HaloTag variants for fluorescence lifetime-based multiplexed sensing and imaging^11^. While HaloTag engineering achieves remarkable lifetime modulation of up to 2.9 ns, protein engineering is usually computationally complex and labor-intensive, requiring extensive trial-and-error and iterative selection. Although nucleic acid engineering can be substantially simpler than protein engineering, there has not been any demonstration of effective nucleic acids engineering to create fluorescent probes with diverse lifetimes for multiplexed detection.

Here we propose a direct, large-scale, rapid, and effective strategy to screen a class of nucleic acid-based probes, termed fluorogenic aptamers^12-15^ (FAPs, such as *Broccoli*^16^, *Corn*^17^, and *Mango*^18^), for diverse fluorescence lifetimes. Fluorogenic aptamers are increasingly important in biosensing and bioimaging^15, 19, 20^, not only due to their compatibility with live-cell applications^21, 22^ but also because they offer excellent signal-to-background ratios in metabolite sensing^21^ and mRNA imaging^22^. Upon binding their cognate small-molecule fluorogens, FAPs often stabilize the planar structures of these fluorogens, suppressing non-radiative decay pathways and promoting radiative decay, which results in marked fluorescence enhancement^23^. Besides, FAPs offer high structural programmability and undergo large conformational changes upon binding^24, 25^, making them suitable for designing sensors with enhanced sensitivity^26, 27^.

Recent developments have led to the creation of several RNA-based fluorogenic aptamers (RNA-FAPs) for molecular sensing, including *Broccoli*/DFHBI-1T for 5-hydroxytryptophan (5-HTP) recognition^28^ and *Mango*/TO1-biotin for small non-coding RNA imaging^19^. Although RNA-FAPs of different emission colors have been developed^15^ and used for multiplexed detection^29, 30^, cross-reactivity with fluorogens and overlapping emission spectra substantially complicate color-based multiplexing and can compromise detection fidelity^15, 31, 32^. Moreover, RNA-based aptamers are inherently susceptible to RNase-mediated degradation in biological environments, limiting their robustness for imaging and quantitative applications^21, 33^.

These limitations can be overcome by developing a set of DNA-based fluorogenic aptamers (DNA-FAPs) that bind the same fluorogen but exhibit distinct fluorescence lifetimes. Single-stranded DNA can be expressed in live cells using retron-based^34, 35^ or reverse transcriptase-based^36^ systems, and the intracellular stability of DNA aptamers can be further enhanced by co-expression of the β protein^36^. Since all DNA-FAP variants within a given set bind the same fluorogen, cross-reactivity is inherently eliminated in multiplexed detection schemes. Although DNA-FAPs, such as *DIR aptamers*^36, 37^ and *Lettuce*^38, 39^, have been demonstrated for pathogenic nucleic acid detection *in vitro*^39^ and metabolite sensing in live cells^36^, no emission color palette exists for DNA-FAPs, let alone a set of DNA-FAP variants with diverse fluorescence lifetimes.

While crystal structures of several FAPs^38, 40^ have been solved, their utility in guiding rational design of new FAP variants with tailored photophysical properties remains limited. One notable exception is the work from Jaffrey’s group, which achieved a 20-nm emission spectral tuning range of DFHO complexed with *Broccoli* variants differing by only a single nucleotide^41^. Inspired by this result, we hypothesize that similar mutational strategies can enable precise control over other photophysical properties of FAPs, such as fluorescence lifetime. Rather than relying on structure-guided rational design, we demonstrate that direct large-scale screening provides a more effective route for diversifying FAPs toward new functional outcomes. Importantly, the resulting screening data also yield valuable structural and functional insights into aptamer-fluorogen complexes, creating a feedback loop for further optimization^4^.

Our method relies on a repurposed next-generation sequencing (NGS) platform and fluorescence lifetime imaging microscopy (FLIM) characterization, which we term FAP-FLIM-NGS (fluorogenic <u>ap</u>tamer-based fluorescence lifetime imaging <u>m</u>icroscopy on <u>n</u>ext-<u>g</u>eneration <u>s</u>equencing chips), to rapidly screen ∼10^4^ FAP variants and identify those that, when complexed with their designated fluorogens, exhibit substantially distinct fluorescence lifetimes suitable for multiplexed detection. We have previously adopted a similar repurposed next-generation sequencing platform termed CHAMP^42^ (Chip-Hybridized Association-Mapping Platform) to screen ∼40,000 fluorescent nanomaterial species on Illumina *MiSeq* flow cells and identified those with bright emission and distinct colors^43^. Building on this prior work, here we prepared a library of ∼10^4^ *Lettuce* variant strands, introduced them into a *MiSeq* flow cell, and sequenced them. Then we conducted fluorescence lifetime screening of these *Lettuce* variants after complexing them with the fluorogen TO1-biotin. From the successful lifetime measurements on 8,821 variants, three *Lettuce* mutants exhibiting markedly different fluorescence lifetimes were identified: a single mutant (smC14T, 6.0 ns) and two double mutants (dmA5T/C14T, 5.2 ns; dmA5T/T22A, 4.4 ns). Our FAP-FLIM-NGS pipeline enables the identification of functionally distinct FAP variants from a library of ∼10^4^ strands within a single week (from library design and sequencing to lifetime imaging and data mapping), representing a major advancement over traditional aptamer selection and optimization methodologies such as SELEX^44^ and microfludics^19^.

To validate the utility of the selected *Lettuce* variants, a set of split *Lettuce* sensors^39, 45^ (hereafter denoted as splitLet) were prepared to target thee pathogenic nucleic acids. In this design, each splitLet pair comprised split “signaling fragments” derived from an identified *Lettuce* variant and “targeting fragments” that bind the target strand at adjacent sites, in a typical binary probe^46^ arrangement. Upon hybridization with a target sequence, the signaling fragments reassembled into a functional *Lettuce* core, enabling incorporation of TO1-biotin and resulting in strong fluorescence emission with a characteristic lifetime. By systematically optimizing stem configurations and magnesium ion concentrations, we achieved highly specific detection of target RNAs. Using a phasor analysis^47^, we demonstrated simultaneous detection of three viral RNA markers, SARS-CoV-2, MERS-CoV, and Influenza A, in a single reaction. Each viral target produced a distinct fluorescence lifetime signature, enabling accurate discrimination of the three RNA species based on their respective lifetimes.

Beyond this proof-of-concept, the FAP-FLIM-NGS pipeline represents a transformative improvement in fluorogenic aptamer screening, offering an intensity-independent, high-throughput method for systematic optimization of aptamer/fluorophore pairs. By enabling the selection of FAPs with distinct fluorescence lifetimes, the FAP-FLIM-NGS pipeline significantly expands the potential of FAP-based multiplexing. These findings lay the groundwork for developing next-generation molecular tools designed to meet the increasing demand for sensitive, specific, and multiplexed detection in both research laboratories and clinical settings.

## RESULTS

### FAP-FLIM-NGS pipeline: high-throughput selection of FAPs using FLIM on NGS chips

In our recent studies, we noticed that *Lettuce*/TO1-biotin complex is at least four-fold brighter than the original *Lettuce*/DFHBI-1T conjugate^4^. This enhanced brightness greatly facilitated FAP-FLIM-NGS screening aimed at discovering aptamer variants with novel photophysical properties. For systematic analysis, positions within the original *Lettuce* aptamer design were categorized into four structural regions: P1 stem, upstream binding core, downstream binding core, and P2 hairpin (**Figure 1a**).

**Figure 1.**
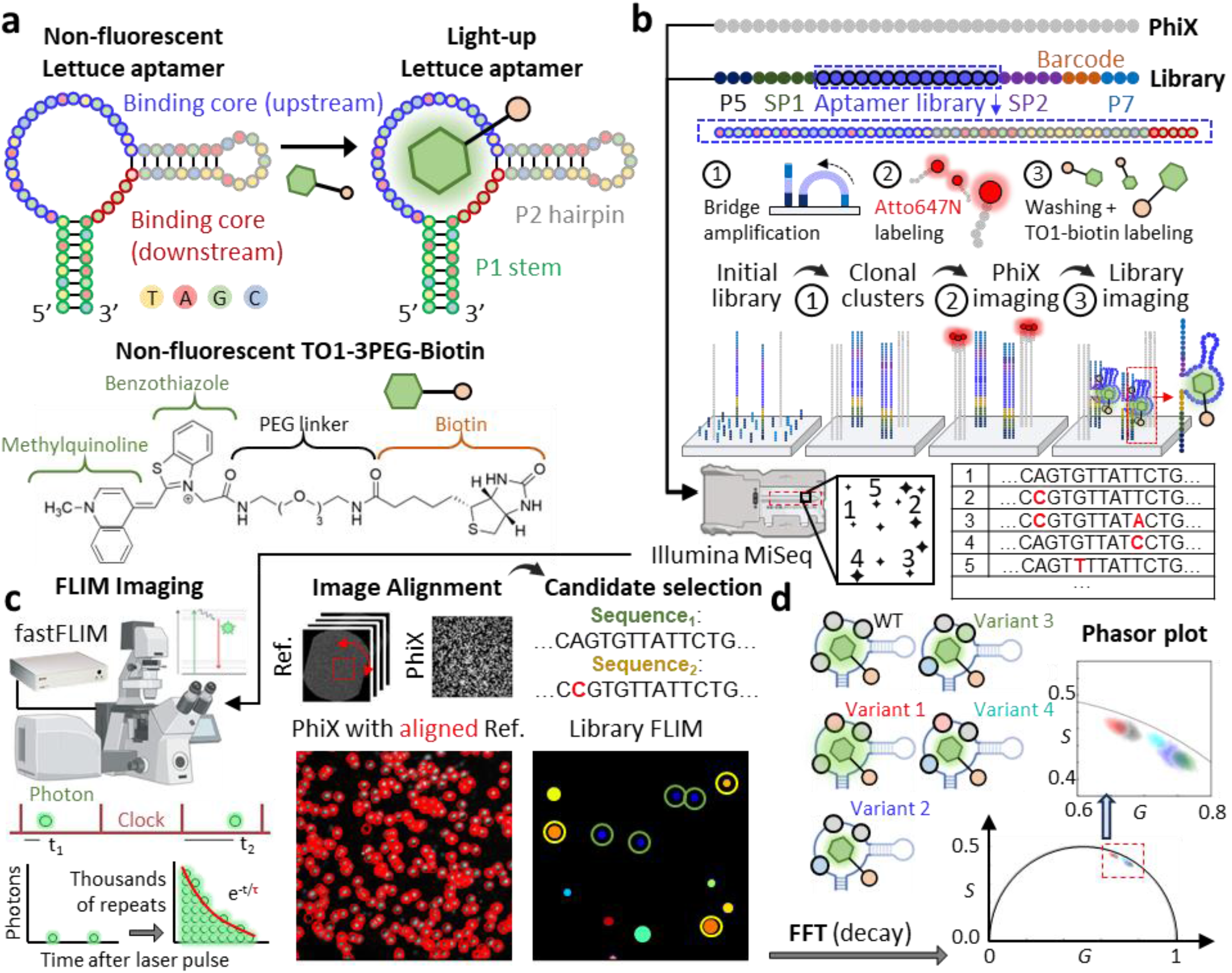
Multiplexed RNA detection using the modular FAP-FLIM-NGS screening platform. (**a**) A schematic shows the structural design of the Lettuce DNA aptamer, which consists of four key regions: the P1 stem (green-bordered), upstream binding core (blue-bordered), downstream binding core (red-bordered), and P2 hairpin (gray-bordered). The aptamer selectively binds the fluorogenic dye, TO1-3PEG-Biotin, inducing a fluorescence signal upon activation. (**b**) The aptamer library is inserted between sequencing primers SP1 and SP2, flanked by P5 and P7. Both the PhiX control and the aptamer library are loaded onto the *MiSeq* chip, followed by bridge amplification to generate clonal clusters. Initial imaging is performed using Atto647N-labeled PhiX cDNA to determine reference cluster positions. This is followed by TO1-biotin labeling to visualize the aptamer library via fluorescence. (**c**) After fluorescence lifetime imaging using the fastFLIM system, PhiX clusters serve as spatial references for image alignment, enabling accurate mapping and analysis of individual aptamer clusters. Aptamer sequences are then matched to their corresponding fluorescence lifetime measurements to determine the lifetime profiles of individual variants. **(d)** The resulting FLIM data are processed using a fast Fourier transform (FFT) to convert pixel-wise fluorescence decay signals into phasor plots. These plots provide a visual fingerprint of fluorescence lifetime characteristics for each aptamer candidate. The presence of distinct phasor cluster patterns among variants highlights the ability of the FAP-FLIM-NGS platform to discriminate multiple aptamer species in parallel based on their unique lifetime profiles.

To tune the fluorescence lifetime of the canonical *Lettuce*/TO1-biotin complex (τ = 5.8 ns to start with), we generated a library of ∼10^4^ *Lettuce* variants covering single mutations (sm), double mutations (dm), single or double insertions (si or di), and single or double deletions (sd or dd) at defined positions (**Figure 1b** and **Supplementary Methods**). The library was synthesized and loaded onto an Illumina *MiSeq* flow cell following standard sequencing protocols, with PhiX fiducial markers included to facilitate spatial alignment between sequencing data and fluorescence images^43^. After sequencing, Atto647N-labeled probe and TO1-biotin were introduced to the flow cell to light up fiducial markers and *Lettuce* variants, respectively (**Supplementary Methods**), and the flow cell was scanned using a confocal FLIM system^48^ in two separate excitation/emission channels (**Figure 1c** and **Supplementary Figure S1**). After collecting digital-frequency-domain fluorescence response data at each pixel, FLIM images were generated using lifetime values obtained through nonlinear least-squares fitting of the multifrequency modulation and phase data^47, 49^. By employing an in-house alignment algorithm to map fiducial markers in the FLIM images to the sequencing FASTQ data, both the average fluorescence intensity and the lifetime of each *Lettuce*/TO1-biotin variant in the library could be determined (**Supplementary Figure S2** and **Supplementary Methods**).

FLIM screening revealed *Lettuce* variants exhibiting distinct fluorescence lifetimes, reflecting their differences in binding TO1-biotin and activating its fluorescence. At least 5 distinguishable clusters were observed in the phasor plot, which not only underscore variations in fluorescence decay dynamics, but also demonstrate that FAP-FLIM-NGS is an effective way to rapidly identify variants with similar photophysical behaviors (**Figure 1d** and **Supplementary Figure S3**). Each cluster represented a variant and was constructed from decay histograms aggregated over all pixels corresponding to that variant. By selecting representative variants that occupy well-separated clusters, a set of aptamers could be formed to attain lifetime-based multiplexed detection of a set of pathogenic nucleic acids.

### Characterization of changes in photophysical properties *via* mutagenesis

To assess how specific mutations alter photophysical properties of the *Lettuce*/TO1-biotin complex, we systematically analyzed single and double mutants, as well as insertion and deletion variants. Heatmaps were used to visualize changes in fluorescence lifetime and intensity relative to the wild-type canonical *Lettuce* (WT; **Figure 2a, b**; **Supplementary Figures S4–7**). To ensure data quality, variants represented by fewer than five polony replicates in intensity images were excluded from further analysis (**Figure 2a, b** and **Supplementary Figures S4-5**). Although many variants showed reduced fluorescence intensity, changes in fluorescence lifetime followed distinct patterns, consistent with the fact that quantum yield and lifetime are not always directly correlated. Specifically, while 72% of single mutants and 94% of double mutants showed reduced intensity, only 64% of single mutants and 78% of double mutants displayed shorter lifetimes (as compared to the WT *Lettuce*; **Figure 2a, b**). This divergence demonstrates that fluorescence lifetime could provide information beyond intensity, offering insights into the molecular organization surrounding the fluorogen within the FAP/fluorogen complex.

**Figure 2.**
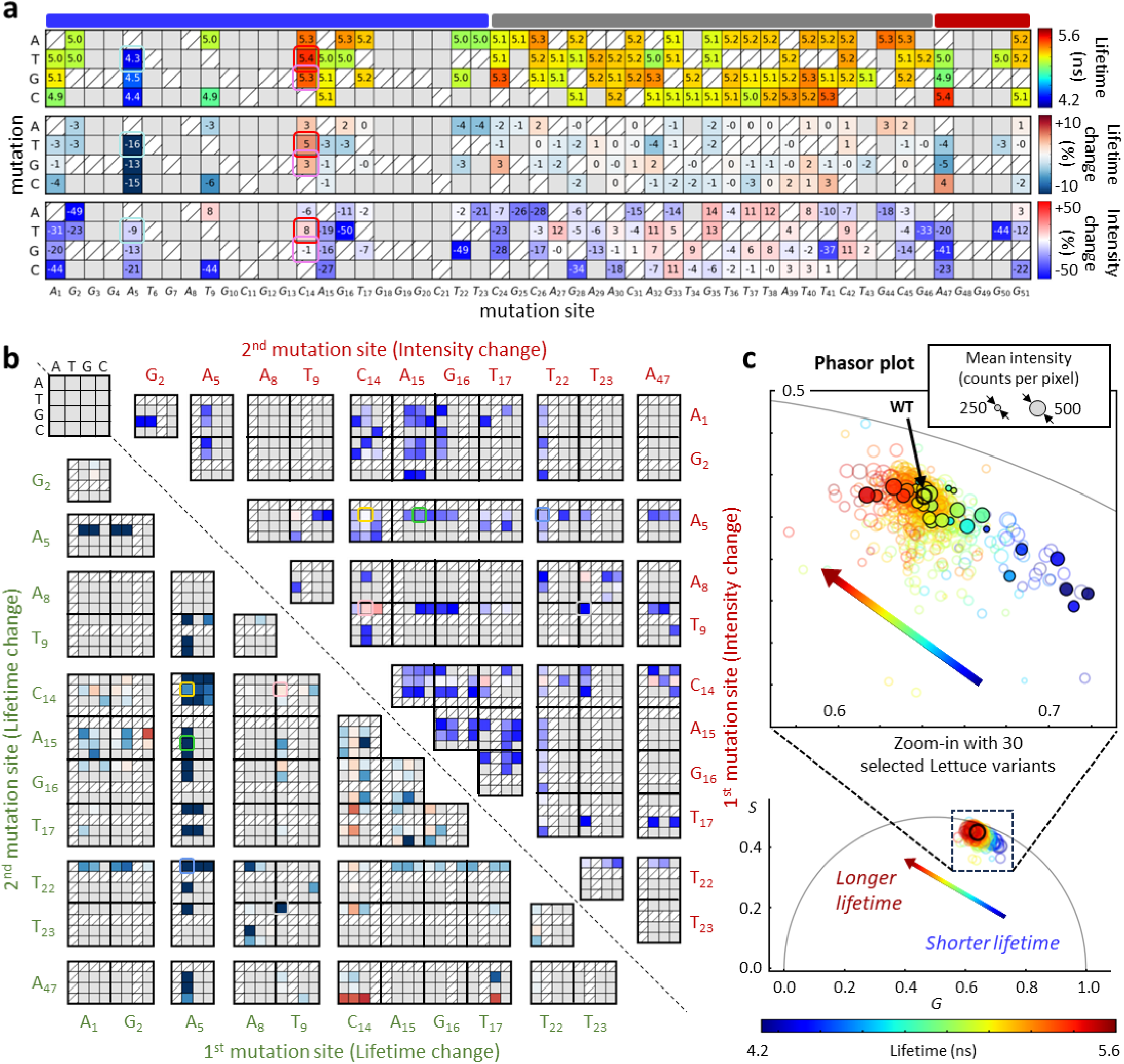
Systematic characterization of fluorescence lifetime and intensity changes in Lettuce variants. (**a**) Heatmaps summarizing the effects of single-nucleotide mutations across the Lettuce aptamer sequence. The top panel displays mean fluorescence lifetime values on chip, the middle panel shows percentage changes in lifetime relative to the wild-type (WT) sequence, and the bottom panel reports percentage changes in fluorescence intensity. Mutations are arranged by position (columns) and substituted base (rows). Gray shading denotes variants with insufficient data (fewer than five polony replicates). Colored blocks above the heatmaps indicate different regions of the sequence: blue and red for the upstream and downstream chromophore-binding cores, gray for the P2 hairpin. (**b**) 2D matrix heatmaps showing pairwise effects of double mutations. The lower triangle represents percentage changes in fluorescence lifetime, while the upper triangle indicates corresponding changes in fluorescence intensity. (**c**) Phasor plot mapped all analyzed Lettuce variants. Fluorescence decay histograms from individual pixels within each polony were subjected to fast Fourier transform (FFT) to generate phasor coordinates. Each variant is represented as a single circle, where the position corresponds to the mean Fourier-space coordinates averaged over all pixels associated with that variant and the circle size reflects fluorescence intensity (photon counts per pixel). Color indicates the mean lifetime value according to the colorbar. The black circle represents the WT, while solid-filled circles indicate 30 variants selected for in-solution validation. The zoom-in panel highlights a subset of 30 candidates with well-resolved lifetime profiles spanning from shorter to longer lifetimes, supporting their utility for multiplexed sensing applications.

For single mutants, heatmaps of lifetime values and percentage changes revealed clear position-dependent effects (**Figure 2a**). Mutations at position 5 (A→T, G, or C) caused the largest reductions in fluorescence lifetime (−13% to −15%), whereas mutations at position 14 (C→A, T, or G) consistently increased lifetime (+3% to +5%). Mutations at position 47, however, showed diverse outcomes: an A→C substitute increased lifetime by 4%, while A→T and A→G substitutes decreased lifetime by 4% and 5%, respectively, underscoring the site’s context-dependent behavior. For double mutants, a composite heatmap was used to compare lifetime percentage changes (lower triangle) alongside intensity percentage changes (upper triangle; **Figure 2b**). Compared to intensity readouts, fluorescence lifetime offered greater discriminatory power among variants, thereby enhancing multiplexing potential.

In the double mutation heatmap, distinguishing intensity changes between variants is challenging. However, fluorescence lifetime provides clearer differentiation, particularly between variants with a mutation at position 5 and those without. Variants with a mutation at position 5 predominantly exhibit fluorescence lifetime reductions greater than 10%, whereas those without the mutation generally show only minor decreases or even increases exceeding 10%. Conversely, mutations at position 14 generally increased lifetime across various combinations, even in the presence of secondary mutations. Interestingly, variants with combined mutations at positions 5 and 14 still exhibited shorter lifetimes, suggesting that the mutation at position 5 may have a dominant influence on photophysical properties.

Phasor analysis^9, 47, 48^ was employed to characterize the fluorescence lifetimes of all *Lettuce* variants within the library. Photon arrival time histograms were collected at the pixel level within each sequencing polony (∼15 pixels in size; **Supplementary Figure S8**) and processed using fast Fourier transform (FFT) to convert the decay histogram of each pixel into Fourier space^50^ (**Figure 2c** and **Supplementary Methods**). Each variant was represented as a single circle in the phasor plot, with its position defined by the mean Fourier-space coordinates averaged over all pixels associated with that variant. The circle color and size encoded the mean fluorescence lifetime and intensity, respectively, calculated on a polony-wise basis. This dual-parameter visualization scheme enabled efficient assessment of both lifetime separability and signal robustness. Variants exhibiting distinct clustering with minimal overlap in phasor space were prioritized, as lifetime separability in phasor space directly reflects their potential for lifetime-based multiplexing. Fluorescence intensity served as an additional filtering criterion to exclude dim variants which could not provide sufficient signal-to-noise ratios for sensing. Using these criteria, 30 *Lettuce* variants with favorable photophysical properties were selected for subsequent in-solution validation (**Supplementary Table S1**).

### In-solution validation of chip-selected FAPs with diverse lifetimes

To validate the reliability of on-chip screening, 30 *Lettuce* mutants were selected for in-solution validation, including 10 single mutations, 11 double mutations, 8 single insertions, and 1 single deletion (**Figure 3a** and **Supplementary Table S1**). Although mean fluorescence lifetimes measured on the *MiSeq* chip were inherently shortened due to higher background noise, chip- and solution-based results were in strong agreement, exhibiting highly consistent percentage changes in fluorescence lifetime (R^2^ = 0.87). For example, the smA5T mutant showed a 16% reduction in fluorescence lifetime on the *MiSeq* chip and an 18% reduction in solution, while smC14T exhibited identical lifetime increases of 5% across both platforms.

**Figure 3.**
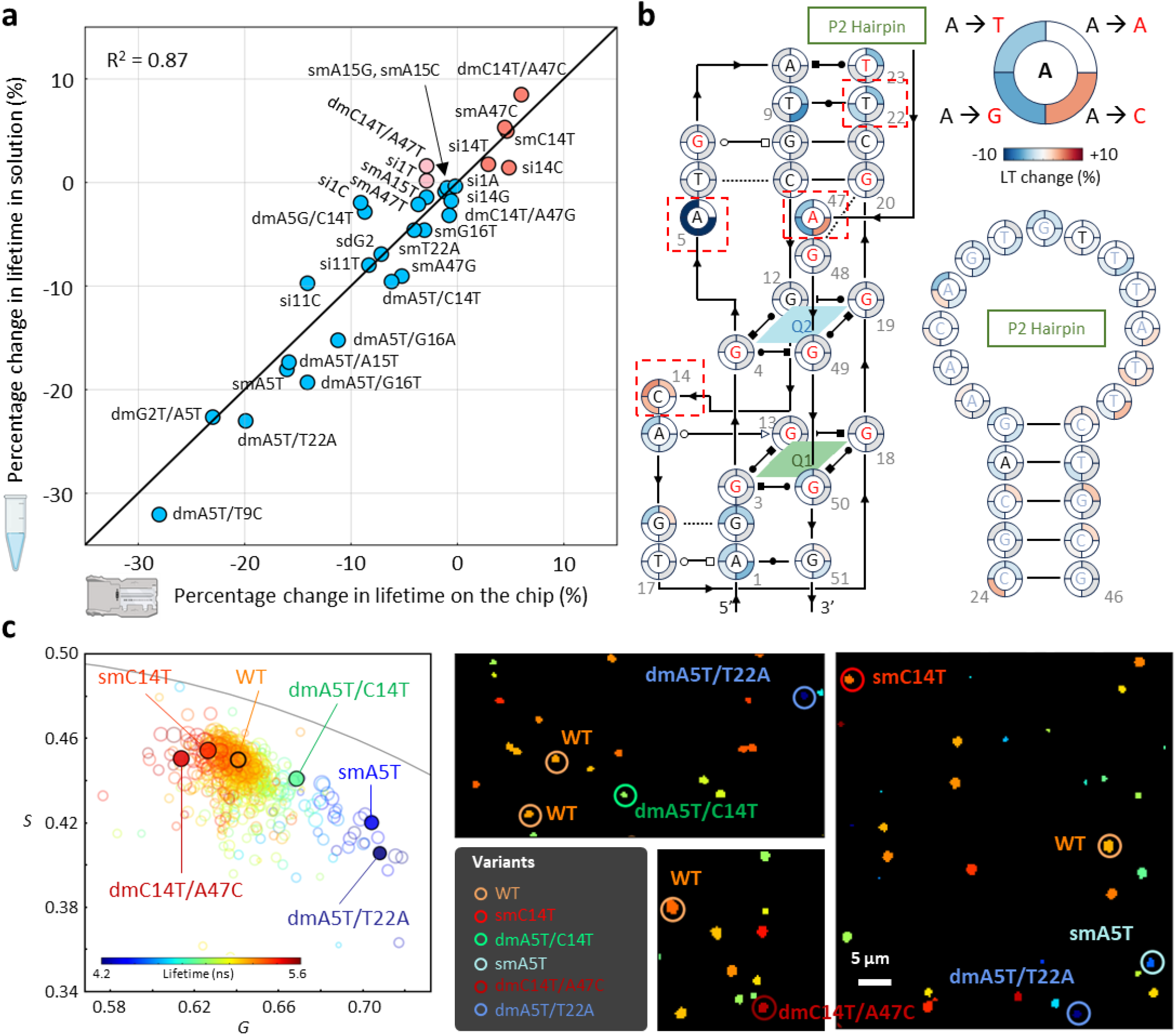
In-solution validation of chip-based FAP-FLIM-NGS screening results on Lettuce variants. (**a**) Correlation analysis of fluorescence lifetime changes measured on-chip versus in-solution for a panel of 30 Lettuce variants. The scatter plot compares the percentage change in mean fluorescence lifetime for each variant relative to the wild-type aptamer, with on-chip measurements plotted on the x-axis and in-solution values on the y-axis. The strong linear correlation (R^2^ = 0.87) confirms the reproducibility and robustness of chip-based screening. (**b**) Structural mapping of the single-nucleotide mutations onto the predicted secondary structure of Lettuce. Red boxes indicate positions with mutations tested in panel (**a**), while the donut plot next to each base illustrates the magnitude of lifetime change for each type of nucleotide substitution. The structure includes two G-quadruplex motifs (Q1 and Q2) and a P2 hairpin, with mutations at positions 5, 14, 22, and 47 showing the significant impact on lifetime modulation. (**c**) Phasor-based visualization and representative FLIM images of selected variants. The left panel displays a phasor plot of fluorescence lifetime signatures for canonical Lettuce (WT, orange) and engineered variants (smA5T, cyan; smC14T, red; dmA5T/C14T, green; dmA5T/T22A, blue; dmC14T/A47C, dark red). Right panels show corresponding FLIM images, with individual polonies color-coded by variant. Scale bar, 5 µm.

Mapping the tested mutations onto the secondary structure of the *Lettuce* aptamer highlighted key nucleotide positions implicated in photophysical modulation^38^ (**Figure 3b, red dashed box**). Among the 30 mutants tested, positions 5, 14, 22, and 47 were identified as critical sites for fluorescence lifetime modulation. Mutations at position 5 consistently decreased fluorescence lifetime across all variants, with reductions of up to 18%, whereas mutations at position 14 uniformly increased fluorescence lifetime by up to 5%. Consistent with the on-chip results, mutations at position 47 produced diverse outcomes in solution. Position 22 also contributed to lifetime modulation, particularly when combined with other mutations.

Mutational perturbations within the P2 hairpin region (nucleotides 24-46) generally produced milder and more uniform effects on fluorescence lifetime than those in the fluorogen-binding core (**Figure 3b**). However, structural alterations in this region, such as insertions or deletions, were frequently accompanied by a marked reduction in fluorescence intensity (**Figure 2a** and **Supplementary Figures S6–S7**). This finding suggested that although the P2 hairpin is not a primary determinant of lifetime modulation, its structural integrity is essential for efficient activation of TO1-biotin. Together, these results indicate that the fluorogen-binding core is the dominant contributor to lifetime tuning, while the P2 hairpin indirectly influences sensing performance by modulating signal strength.

These variants demonstrated cross-platform consistency and showed clear separability in both phasor space and FLIM images (**Figure 2c** and **Figure 3c**). Collectively, these findings validated the FAP-FLIM-NGS pipeline as a reliable approach for high-throughput characterization of FAPs. The in-solution measurements further confirmed the reproducibility of lifetime modulation and highlighted the potential of FAPs for lifetime-based multiplexed sensing.

### Design of split *Lettuce* sensors for RNA detection

To explore the applicability of *Lettuce* variants for RNA detection, we drew inspiration from the work of Jaffrey’s group^39^ to engineer split *Lettuce* (splitLet) pairs that reconstitute into a functional fluorogen-binding core upon hybridization to a target RNA (**Figure 4a**). Each splitLet probe consists of two components and is non-fluorescent on its own. Short flanking sequences are appended to each component to enable specific target recognition. Fluorescence is activated when the two components are brought into proximity through hybridization in a juxtaposition to a synthetic target RNA, facilitating reconstruction of the functional *Lettuce* core.

**Figure 4.**
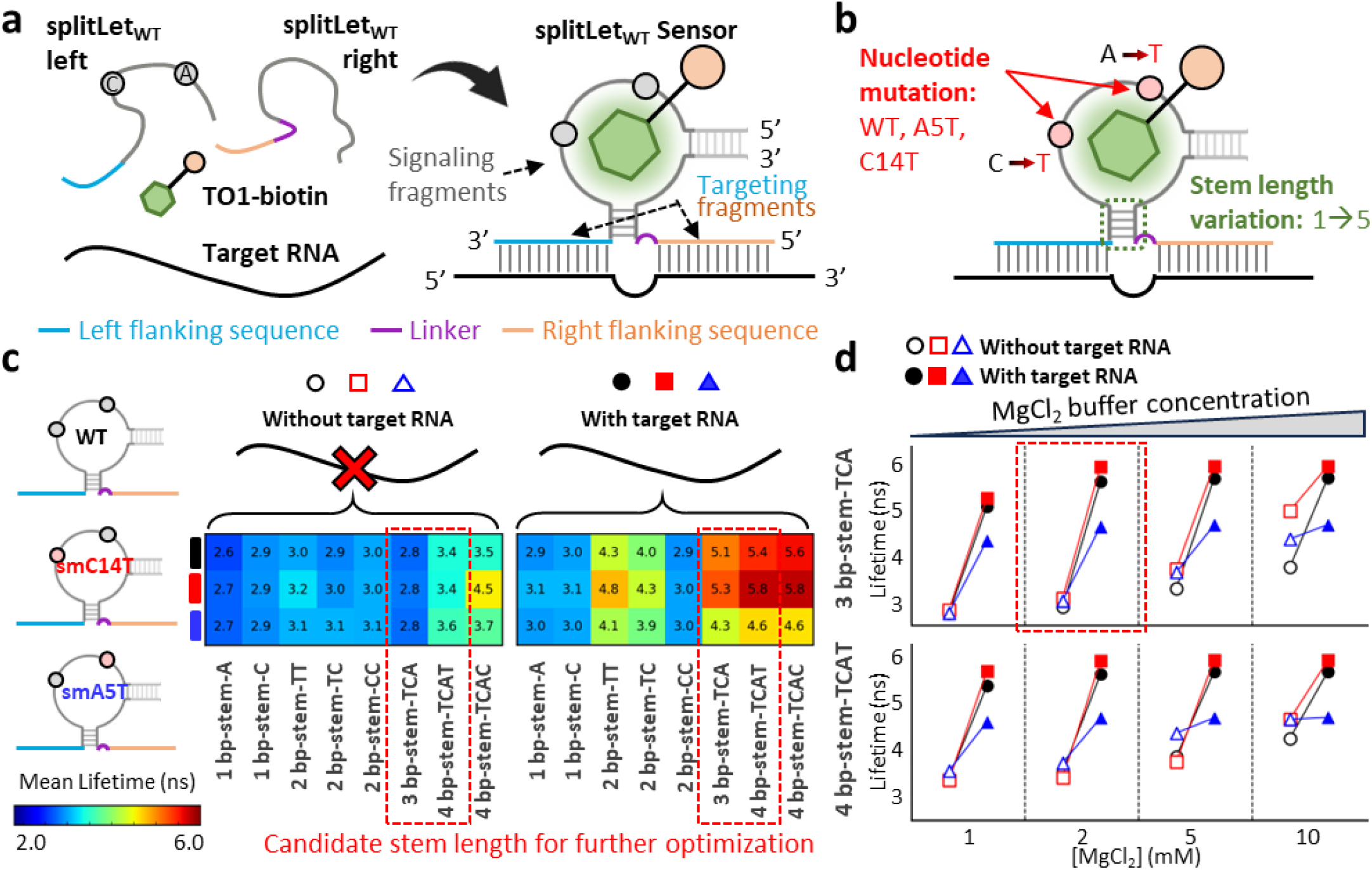
Optimization of splitLet sensors for RNA detection. This figure presents the design and systematic optimization of splitLet RNA sensors for the selective detection of target RNA sequences. (**a**) A schematic illustrates how two splitLet DNA fragments, each carrying complementary flanking sequences (blue and orange), assemble into a functional fluorescent sensor in the presence of a specific target RNA, forming a three-way junction. The binding of TO1-biotin activates fluorescence. (**b**) Variants of the sensor were engineered by introducing point mutations (WT, smA5T, smC14T), altering stem lengths (1–5 base pairs) and sequence compositions to assess performance. (**c**) A heatmap compares fluorescence lifetimes across different mutation-stem length combinations, both in the absence and presence of target RNA. Optimal candidates (highlighted in red boxes) show significant lifetime shifts upon target binding and were selected for further evaluation. (**d**) Line plots display fluorescence lifetime responses of the top sensor candidates (WT: black, smA5T: blue, smC14T: red) under increasing MgCl^2^ concentrations, highlighting the importance of buffer conditions in modulating sensor performance.

To optimize the splitLet system for discriminating between target and non-target conditions, we systematically designed constructs with varying P2 stem lengths (ranging from 1 to 5 stems) and GC contents (**Figure 4b, c; Supplementary Table S2**). Variations in stem length and GC content are critical for fine-tuning binding core reassembly, enabling efficient TO1-biotin binding and fluorescence activation with characteristic lifetimes in the presence of target RNA. We hypothesized that if the P2 stem is short and unstable, effective core formation does not occur even in the presence of target RNA, leaving splitLet inactive. Conversely, if the P2 stem is long or overly stable, spontaneous core formation can occur in the absence of target, rendering non-specific activation of splitLet. Here we aimed to identify a P2 stem design that not only yields the brightest emission but also gives full recovery of characteristic fluorescence lifetime upon addition of target RNA to the solution.

To refine the system further, we incorporated nucleotide mutations and examined WT Lettuce alongside the smC14T and smA5T variants as representative examples to demonstrate effects on fluorescence lifetime (**Figure 4c; Supplementary Figure S9; Supplementary Table S2**). In the absence of target RNA, fluorescence lifetimes of all splitLet constructs were consistently shortened, ranging from 2.6 ns to 3.7 ns across various P2 stem designs, with the exception of the 4 bp-stem-TCAC splitLet_smC14T design (**Figure 4c**). For P2 stem lengths shorter than 3 bp, fluorescence lifetimes remained below 3.1 ns, suggesting limited core reconstitution without a target. In contrast, for P2 stem lengths longer than 3 bp, fluorescence lifetimes became longer and exceeded 3.4 ns, indicating partial stabilization of the binding core even in the absence of target RNA. Upon addition of target RNA, fluorescence lifetimes increased significantly when P2 stem length exceeded 2 bp. Based on these results, the 3 bp-stem-TCA and 4 bp-stem-TCAT configurations were identified as the most effective for differentiating target and non-target conditions. The largest lifetime shifts were observed for the 3 bp-stem-TCA designs, with fluorescence lifetimes increasing by 82% for splitLet_WT, 89% for splitLet_smC14T, and 53% for splitLet_smA5T, indicating strong target binding and efficient core reconstitution. Although the 4 bp-stem-TCAT designs also produced substantial lifetime shifts upon target binding, the contrast between target and non-target conditions was less pronounced compared to the 3 bp-stem-TCA designs.

We next optimized magnesium ion (Mg^2+^) concentration to enhance splitLet sensor performance and stability (**Figure 4d**). Compared with in-solution lifetime measurements of intact *Lettuce* variants (**Figure 3a**), in which smC14T and smA5T complexed with TO1-biotin exhibited lifetimes of 6.0 ns and 4.7 ns, respectively, lifetimes of splitLet constructs measured at 1 mM MgCl^2^ in the presence of target were shorter. This result suggested that low Mg^2+^ concentrations provide insufficient stabilization of splitLet, limiting binding core reconstitution. At 2 mM MgCl^2^, fluorescence lifetimes of splitLet sensors with targets closely matched those of the intact *Lettuce* variants, underscoring the essential role of Mg^2+^ in stabilizing the splitLet and enabling proper core formation. At high MgCl^2^ concentrations (5 mM and 10 mM), however, lifetime shifts between target and non-target conditions diminished, likely due to overstabilization of the splitLet and partial core formation in the absence of target RNA. Together, these results identified the 3 bp-stem-TCA P2 configuration and 2 mM MgCl^2^ as optimal parameters for splitLet-based, lifetime-resolved RNA detection.

### Validation of splitLet’s specificity in viral RNA detection

Based on combined intensity and lifetime performance relative to wild-type *Lettuce*, three variants (smC14T, dmA5T/C14T, and dmA5T/T22A) were selected as top candidates for lifetime-based multiplexed sensing, with smA5T included for additional validation of measurement consistency (**Supplementary Figure S10**). To evaluate the specificity of splitLet sensors in viral RNA detection, we prepared RNA targets corresponding to markers from human-infective viruses associated with severe diseases, including SARS-CoV-2, MERS-CoV, and Influenza A^51, 52^ (**Figure 5** and **Supplementary Table S3**). Multiple splitLet sensors were designed with optimized left and right flanking sequences targeting specific regions of each of these viral RNAs. Sensors bearing the four respective variants were individually tested against each RNA target to confirm detection specificity. To further determine whether each sensor could selectively bind its cognate target under competitive conditions, all three viral RNA targets were combined in a single tube, and only one specific sensor was introduced at a time. In total, five experimental conditions were evaluated for each sensor: sensor only, three single-target conditions (SARS-CoV-2 RNA, MERS-CoV RNA, or Influenza A RNA), and one mixed-target condition containing all three RNAs. Because each sensor incorporated four variants, this corresponded to 20 measurements per sensor. Across the three sensors, a total of 60 sensor–target combinations were evaluated.

**Figure 5.**
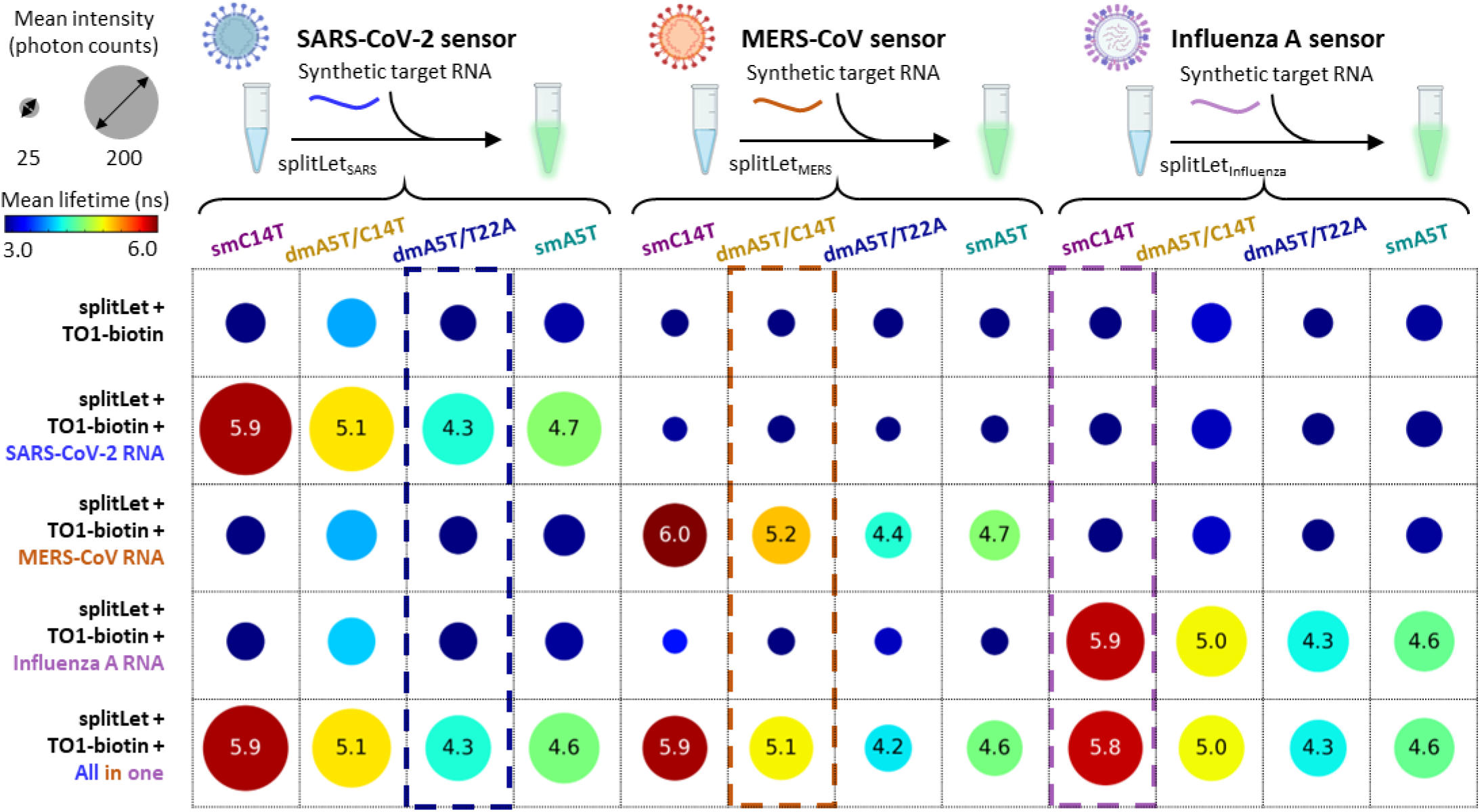
Specificity of three splitLet sensors in detecting synthetic SARS-CoV-2, MERS-CoV, and Influenza A viral RNA targets, respectively. A total of 60 sensor–target combinations were tested, comprising three synthetic viral RNA targets and splitLet sensors with four respective variants FAP constructs per target, tested across five experimental conditions. Each column represents a specific variant, smC14T (purple), dmA5T/C14T (brown), dmA5T/T22A (blue), and smA5T (green), within the three categories of splitLet sensors (splitLet_SARS_, splitLet_MERS_, splitLet_Influ_), within the three splitLet sensor categories (splitLet_SARS_, splitLet_MERS_, splitLet_Influ_), while each row corresponds to a specfic mixing condition. The values within each circle indicate the mean fluorescence lifetime (in nanoseconds), and the circle size reflects fluorescence intensity (average photon counts). In the “All-in-one” condition, all three synthetic RNAs were present simultaneously.

In the presence of the cognate target RNA, we observed at least a 2.5-fold increase in fluorescence intensity, accompanied by a consistent shift in fluorescence lifetime toward the characteristic value of each activated splitLet variant. For instance, sensors targeting SARS-CoV-2 displayed lifetimes of 5.9 ns (splitLet_smC14T), 5.1 ns (splitLet_dmA5T/C14T), and 4.3 ns (splitLet_dmA5T/T22A) when exposed to SARS-CoV-2 RNA, compared to lifetimes below 3 ns in the absence of the target (**Figure 5**). We also tested an additional variant, splitLet_smA5T, which was expected to exhibit a characteristic lifetime of 4.7 ns based on the systematic characterization in **Figure 2-3**. As anticipated, binding with the SARS-CoV-2 target did result in a lifetime of 4.7 ns. These results were consistently reproduced in “all-in-one” samples containing all viral RNA targets simultaneously, demonstrating the high specificity of splitLet sensors. Comparable specificity and characteristic lifetime shifts were observed for sensors targeting MERS-CoV and Influenza A. Fluorescence lifetime changes were independent of intensity variations, confirming lifetime as a robust, target-specific, and reproducible readout for viral RNA detection.

### Multiplexed detection of viral RNA using splitLet sensors and phasor plot analysis

We next evaluated simultaneous detection of three synthetic RNA targets in a single reaction (**Figure 6a** and **Supplementary Table S4**). For lifetime-based multiplexing, we selected SplitLet sensors with well-separated characteristic fluorescence lifetimes: splitLet_dmA5T/T22A (4.3 ns characteristic lifetime and targeting SARS-CoV-2 RNA), splitLet_dmA5T/C14T (5.1 ns, targeting MERS-CoV RNA), and splitLet_smC14T (5.9 ns, targeting Influenza A RNA) (**Figure 5**). Fluorescence decay data acquired by the fastFLIM module were transformed into the frequency domain and visualized using phasor plot analysis, where each sensor-target pair formed a distinct cluster corresponding to its unique lifetime signature in phasor space. Phasor plot analysis was performed at the second harmonic frequency (40 MHz) to enhance discrimination among targets (**Supplementary Methods**).

**Figure 6.**
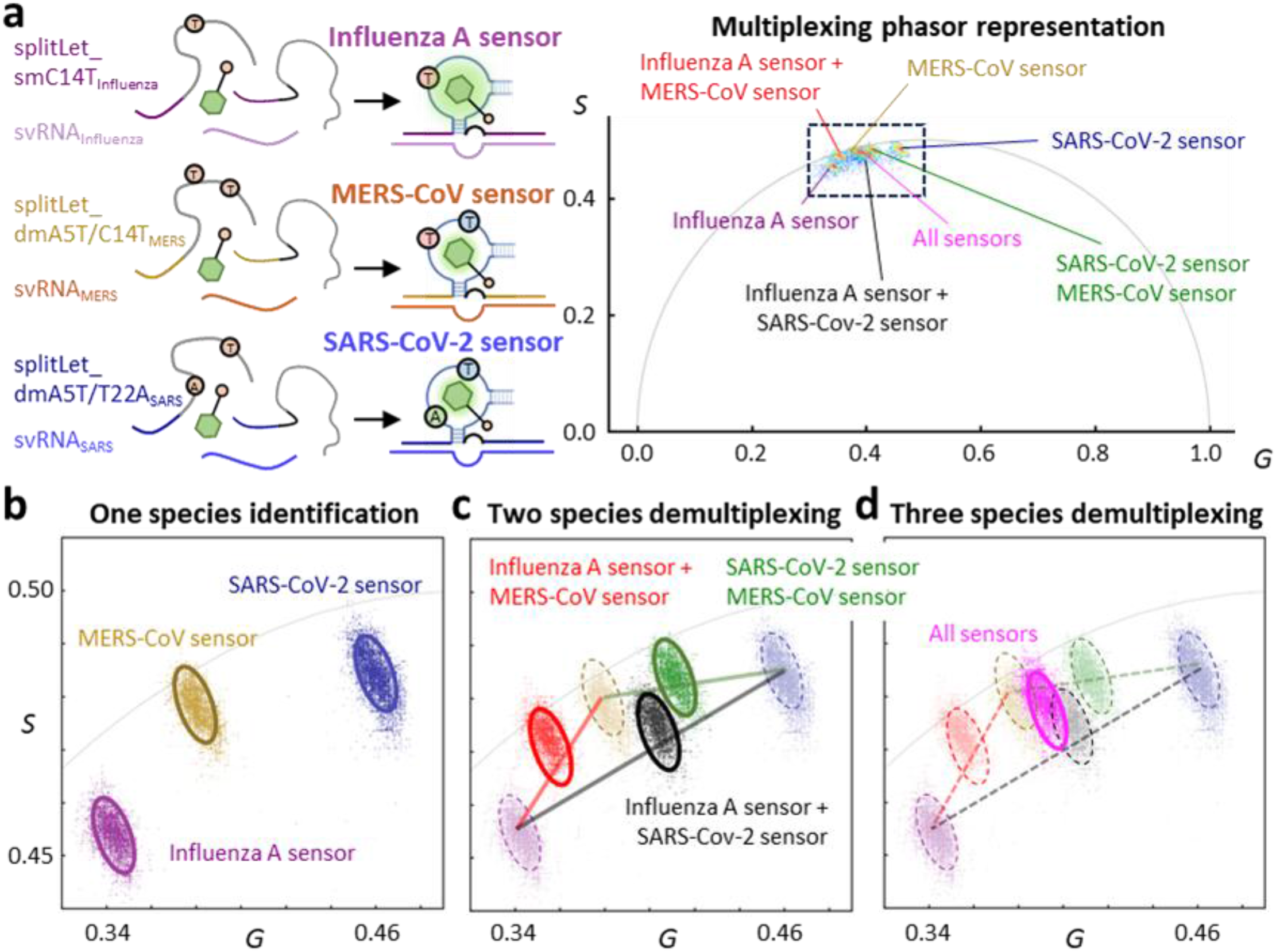
Multiplexed detection of viral RNAs using the phasor FLIM assay. (**a**) Schematics of the splitLet assay design, in which sensors specifically respond to synthetic Influenza A (sRNA_Influenza_), MERS-CoV (sRNA_MERS_), or SARS-CoV-2 (sRNA_SARS_) RNA by reconstituting a fluorescent aptamer structure upon hybridization. The right panel shows the phasor plot summarizing all multiplexed FLIM measurements, where each RNA-specific sensor generates a distinct fluorescence lifetime signature. (**b**) Phasor plot showing single-species identification, with clear clustering for Influenza A (purple), MERS-CoV (orange), and SARS-CoV-2 (blue). (**c**) Two-species demultiplexing, demonstrating accurate separation of Influenza A + MERS-CoV (red), Influenza A + SARS-CoV-2 (black), and SARS-CoV-2 + MERS-CoV (green) based on their composite phasor positions. (**d**) Three-species demultiplexing with all sensors present reveals distinct phasor clusters corresponding to each viral RNA and their composite mixtures. The combined signal from all three viral RNAs is visualized as a unique magenta cluster, demonstrating the capability of phasor-based FLIM analysis to simultaneously resolve multiple RNA targets within a single FLIM imaging assay.

To rigorously evaluate the multiplexing capability of the splitLet sensors, we first introduced each viral RNA target individually, one at a time, into a mixture containing all three sensors under separate experimental conditions. Upon addition of its cognate RNA, the corresponding activated splitLet sensor produced a well-defined and discrete cluster in phasor space, enabling unambiguous discrimination among the three sensor–target pairs (**Figure 6b**). We subsequently analyzed samples containing pairwise combinations of RNA targets—SARS-CoV-2 + MERS-CoV, SARS-CoV-2 + Influenza A, and MERS-CoV + Influenza A (**Figure 6c**). In each condition, the measured fluorescence lifetimes occupied intermediate phasor positions consistent with the weighted contributions of the two activated splitLet sensors. These data demonstrate robust resolution of any two viral targets within mixed samples. Finally, we evaluated an all-target condition in which all three viral RNA targets were combined with the three splitLet sensors in a single reaction (**Figure 6d**). Phasor analysis revealed a well-resolved cluster that could be decomposed into the individual lifetime components associated with each activated splitLet sensor, enabling quantitative determination of their relative fractional contributions^2, 47^.

Together, these results demonstrate that splitLet sensors integrated with fluorescence lifetime measurement and phasor plot analysis provide a robust and versatile platform for multiplexed RNA detection. The system enables reliable discrimination of single-, dual-, and triple-species RNA mixtures using a single excitation source, a single scan, and a simple optical setup requiring only one filter set. Our lifetime-based multiplexing strategy using FAPs represents a unique advance that extends capabilities beyond conventional intensity- and spectrum-based approaches.

## DISCUSSION

Inspired by the seminal work from Johnsson’s group on engineering HaloTag for lifetime multiplexing^11^, here we engineered the fluorogenic aptamer *Lettuce* for the same purpose. Although the attained lifetime tuning range of our engineered *Lettuce* variants (Δτ up to 2.2 ns, as determined from in-solution validation) is smaller than that of engineered HaloTag variants (Δτ up to 2.9 ns), our study demonstrated, for the first time, that fluorescence lifetime can be systematically tuned in a DNA-based fluorogenic aptamer (DNA-FAP). Braselmann’s group recently reported lifetime multiplexing using engineered RNA-FAP *RiboGlow* complexed with Cbl-Atto590^3^; however, the observed lifetime tuning range was very limited (Δτ ∼ 0.2 ns). This modest modulation likely arose because mutations in the *RiboGlow* system do not directly alter the fluorogen microenvironment, but instead indirectly influence the Atto590 lifetime (e.g., by changing the distance between the Cbl quencher and the Atto590 fluorophore). As a result, the small tuning range made phasor analysis and precise quantification challenging. In contrast, in our *Lettuce*/TO1-biotin system, mutations directly perturbed the fluorogen-binding core, thereby altering the local environment of fluorogen and producing substantially larger lifetime shifts. Notably, without using any computational modeling or rational design, our FAP-FLIM-NGS pipeline was able to attain a 2.2-ns lifetime tuning range via screening of ∼10^4^ *Lettuce* variants complexing with TO1-biotin (**Figure 1-3**). Such a large lifetime tuning range has made the phasor-analysis-based multiplexed detection of three viral RNA targets straightforward and highly accurate. By integrating our high-throughput screening pipeline with structure-informed rational engineering, we anticipate that even greater lifetime tuning ranges can be achieved – an avenue we are actively pursuing.

Given that current methods for predicting nucleic acid three-dimensional structures from sequence^53^ remain significantly less advanced than protein structure prediction from amino acid sequences^54^, direct large-scale screening represents a particularly effective strategy for diversifying FAPs toward desired functional outcomes. Moreover, by simply incorporating an *in-vitro* transcription step into the existing FAP-FLIM-NGS pipeline^55, 56^, this approach can be readily extended to engineer RNA-FAPs, such as *Broccoli*^16^, *Corn*^17^ and *Mango*^18^, enabling systematic diversification of their fluorescence lifetimes (not shown in the Supplementary Information). Although cross-reactivity between FAPs and fluorogens remains a key challenge^15, 31, 32^ (for instance, *Corn* and *Red Broccoli* both bind DFHO as well as DFHBI^17, 41^, while *Lettuce, Mango*, and *Corn* all bind TO1-biotin^4, 31^), recent work has demonstrated the feasibility of engineering orthogonal FAP/fluorogen pairs with minimal cross-reactivity for multicolor imaging and sensing^32, 57-59^. We envision that, in the near future, such orthogonal FAP/fluorogen pairs, combined with engineered lifetime variants, will enable higher-order multiplexed detection (6-plex to 9-plex) using pulsed-interleaved excitation^6^ and spectral FLIM techniques pioneered by Sauer^1^, Gratton^2^ and our group^47^. Given the substantially larger repertoire of RNA-FAPs currently available (e.g., *Peach, Mango, Beetroot, Corn*, and *Pepper*, as summarized in a recent review article by Unrau^15^) compared to DNA-FAPs (e.g., *DIR aptamers*^36, 37^ and *Lettuce*^38, 39^), future multiplexed assays may benefit from the combined use of both DNA- and RNA-FAPs. Such color-lifetime “combinatorial” multiplexing strategies could further expand the dimensionality and flexibility of nucleic acid-based sensing platforms.

DNA aptamers are traditionally identified and evolved through Systematic Evolution of Ligands by EXponential enrichment^44^ (SELEX). While the resulting DNA-FAPs exhibit improved photostability and greater resistance to enzymatic degradation over their RNA counterparts^36, 39^, the SELEX process is labor-intensive and time-consuming, typically requiring more than 10 rounds of selection^60^. Moreover, its capacity for fine optimization is limited, as SELEX is primarily designed for broad selection from a pool of ∼10^15^ random sequences, rather than the detailed characterization within 105-107 structurally similar aptamer variants^61^. Although using a pre-defined library^62^ can accelerate optimization, it still requires at least 3-4 rounds of SELEX process^62^ (which takes approximately two weeks to complete) and tedious counterselection against non-target molecules^61^ to refine aptamers. In contrast, our FAP-FLIM-NGS pipeline enables comprehensive screening of ∼10^4^ aptamer variants within a week (from library design and sequencing to lifetime imaging and data mapping), thus representing a major improvement over the traditional SELEX approach for aptamer evolution. Furthermore, we have previously employed a deep learning-based approach termed *flimGANE*^*9, 48*^ to analyze FLIM data and produced consistent outcomes (**Supplementary Figure S11; Supplementary Methods**). Such a machine-learning-based algorithm can be expanded to investigate mutation-lifetime relationships, thus expediting aptamer optimization and repurposing pipeline (much like what is happening to protein engineering field^63, 64^). We estimated, by integrating these advanced ML/AI techniques, the future FAP-FLIM-NGS pipeline has the potential to complete the same process within 2-3 days.

While our current implementation using a *MiSeq* flow cell allows for the screening of only 10^4^ variants, the screening capacity can be substantially expanded (to 105-107 sequences) by adopting higher-throughput flow cells such as *HiSeq, NextSeq*, or *NovaSeq*, without extending the overall optimization timeline. Moreover, the flow cells can be repeatedly used for fluorescence screening up to 15-20 times without showing any major degradation^42, 43^, and the screening results correlate well with test-tube measurements (this report), demonstrating the high reproducibility and reliability of our FAP-FLIM-NGS method. While thorough characterization of structurally similar FAPs provided by the FAP-FLIM-NGS pipeline can effectively identify variants with novel properties, the current approach depends on pre-designed libraries composed of known or closely related variants, which limits *de-novo* discovery from highly diverse combinatorial pools. Nonetheless, future advancements in library design or integration with alternative NGS chip-based selection strategies could help overcome this constraint.

Fluorescence lifetime multiplexing offers several advantages over traditional color- or intensity-based multiplexing approaches^1-3, 47^, including insensitivity to probe concentration, excitation fluctuations, photobleaching, and other artifacts such as background noise ^65-68^, thus resulting in more robust and reliable readouts, especially in complex environments^68^. In addition, fluorescence lifetime measurements can provide valuable information about the local microenvironment surrounding a fluorophore^47^. A notable example is the use of lifetime in RiboGlow to determine the subcellular localization of mRNA or noncoding RNA (e.g., in nucleus, cytosol, or stress granule) in live cells. Compared with intensity-based measurements, lifetime readouts provide both higher contrast and greater reliability^3, 9, 68^. In the present study, many *Lettuce* variants exhibited reduced fluorescence intensity due to structural perturbations; however, their fluorescence lifetime signatures remained robust and clearly distinguishable (**Figure 3-5**), greatly facilitating multiplexed detection (**Figure 6**). Together, these results highlight the complementary nature of fluorescence lifetime and intensity measurements, with lifetime providing an additional and critical dimension of information for applications requiring high specificity and accuracy.

Our screening revealed the critical role of key nucleotide positions, such as A5, C14, and T22, in modulating the fluorescence lifetime and intensity of *Lettuce*/TO1-biotin complex, possibly through various structural mechanisms (**Supplementary Figure S12**). Although changing fluorogen from DFHBI-1T to TO1-biotin does not disrupt the four-way junction structure surrounding the fluorogen^4^, in contrast to DFHBI-1T coordination^38^, TO1-biotin exhibited stronger π-π stacking interactions with surrounding bases with nearly no hydrogen bonding. This enhanced stacking likely contributed to the higher quantum yield observed in the *Lettuce*/TO1-biotin complex (QY = 0.59 for *Lettuce*/TO1-biotin vs. 0.11 for *Lettuce*/DFHBI-1T)^4^. Although smA5T and smC14T display only minor global deviations from the canonical Lettuce/TO1-biotin structure, their local effects differ substantially. In smA5T, T5 rotates outward by 28°, likely loosening local packing and reducing fluorescence. In contrast, T14 in smC14T rotates by 26° but remains >4 Å from the fluorogen, indicating no direct interaction. Instead, enhanced π–π stacking appears to strengthen fluorogen stabilization (**Supplementary Figure S12**). Since the fluorescence lifetime of the double mutant dmA5T/C14T falls between the single mutants (smA5T and smC14T), we hypothesize that these mutations may have an “offset” effect on each other. A more striking example involves the smT22A mutation, which alone results in a relatively shorter fluorescence lifetime of 5.4 ns. Structural data suggest that smT22A may destabilize local architecture, potentially through altered Hoogsteen base-pairing interactions with T9. When combined with smA5T, the double mutant dmA5T/T22A exhibits a markedly shorter lifetime of 4.3 ns. This suggests an “add-on” effect of the two mutations, where smA5T-induced local disruption and altered Hoogsteen base-pairing interactions between A22 and T9 jointly contribute to the behavior of dmA5T/T22A. Future X-ray crystallography studies could provide further insights into the structural basis of these mutation effects on fluorescence lifetime.

In summary, we introduce FAP-FLIM-NGS, a high-throughput, fluorescence-lifetime-based multiplexed sensing platform using engineered fluorogenic aptamer variants. By evolving aptamers with distinct lifetimes and incorporating them into split Lettuce designs, we achieve robust, single-channel discrimination of multiple viral RNA targets without spectral separation. This approach establishes fluorescence lifetime as a generalizable encoding dimension, enabling precise aptamer performance control and target discrimination. Beyond DNA FAPs for multiplexed viral RNA detection, we envision that FAP-FLIM-NGS could be extended to higher-density sequencing platforms and adapted to RNA-based FAPs, providing a scalable, high-throughput foundation for multiplexed, spectrally efficient diagnostics and next-generation molecular sensing.

## Data availability

All data in this study are available from the corresponding author upon reasonable request.

## Acknowledgements

H.-C. Y. is supported by National Institutes of Health (DA060543) and National Science Foundation (2432379 and 2404334).

Y. L. is supported by National Institutes of Health (GM141931) and National Science Foundation (2235455). P. R. is supported by National Institutes of Health (GM106137), Welch Foundation (F-2120), and the Cancer Prevention and Research Institute of Texas (RP210088). C. M. is supported by National Science Foundation (2107393). Y.-I. C. is supported by the Texas Global Seed Grant at UT Austin.

## Contributions

Y.-I. C., Y.-A. K., and H.-C. Y. conceived and defined the project. Y.-I. C. and H.-C. Y. supervised the study. Y.-I. C. and Y.-A. K. designed and performed the experiments. Y.-I. C., Y.-A. K., and S. Y. revised the CHAMP algorithms. Y.-I. C., Y.-A. K., and Y.-J. C. analyzed the data. Y. H., N. S., E. J. B.-S., and C. M. conducted the X-ray crystallography experiments and analyzed the resulting crystal structures. A.-T. N. and S. K. assisted with manuscript revision. Y. W., Z. Y., Y. L., and P. R. contributed to data interpretation and scientific discussion. Y.-I. C., Y.-A. K., and H.-C. Y. wrote the manuscript with input from all authors.

